# Cascaded Nano-ring Dialyzer for Continuous Regenerative Peritoneal Dialysis

**DOI:** 10.64898/2026.09.14.751343

**Authors:** Wonseok Kim, Kihong Kim, Seongjun Hong, Sunhwa Lee, Hyomin Lee, Dong Ah Shin, Seung Hee Yang, Kyunghee Kim, Jeongeun Lee, Kyoung Jin Lee, Woo Sang Cho, Hajeong Lee, Dong Ki Kim, Hee Chan Kim, Myungjin Seo, Donghyun Kim, Luke P. Lee, Yon Su Kim, Jung Chan Lee, Gun Yong Sung, Sung Jae Kim

## Abstract

End-stage kidney disease (ESKD) requires lifelong kidney replacement therapy, yet conventional hemodialysis and peritoneal dialysis remain limited by intermittent treatment, restricted mobility, and repeated dialysate exchange. Here, we developed a continuous regenerative peritoneal dialysis platform based on a cascaded nano-ring-assisted nanoelectrokinetic dialyzer for portable artificial kidney applications. We first identified key geometric and electrokinetic parameters governing the removal performance of nano-ring meshes, including mesh dimension, nano-ring coating thickness, the number of cascaded nano-rings, and the relative orientation between fluid flow and the applied electric field. We then developed a single cascaded nano-ring dialyzer operating at 1.33 mL/min and scaled the treatment capacity to approximately 10 mL/min through parallel integration of multiple modules. In a canine model, the closed-loop system achieved approximately 10 % reduction in uremic toxin levels in serum during continuous regenerative peritoneal dialysis. We further improved the biocompatibility of the regenerated dialysate by integrating plate-type electrodes, activated carbon, bicarbonate buffering, and UV-C treatment. In an unanesthetized canine model, the improved system sustained continuous toxin removal for 4 hours while major hepatic and inflammatory markers remained within recoverable ranges. These results demonstrate that cascaded nano-rings enable continuous dialysate regeneration with scalable throughput, reduced dependence on fresh dialysate, and *in vivo* control of uremic toxins, providing a technological foundation for portable or wearable regenerative peritoneal dialysis systems.

## INTRODUCTION

End-stage kidney disease (ESKD; historically also referred to as end-stage renal disease, ESRD) is a severe chronic condition characterized by irreversible loss of kidney function and is associated with substantial morbidity, mortality, and socioeconomic burden. The global population requiring kidney replacement therapy (KRT) has continued to increase, with historical estimates indicating an annual growth rate of approximately 6–7 % in the treated ESKD population [1]. A systematic analysis of international renal registry data estimated that approximately 2.6 million people received KRT worldwide in 2010 and projected that this number would more than double to approximately 5.4 million by 2030 [2]. This increasing burden has been attributed to population ageing, the growing prevalence of chronic conditions including diabetes, hypertension, and obesity, and improved access to KRT.

Despite substantial advances in KRT, clinical outcomes for patients with ESKD remain poor. In particular, cardiovascular mortality among patients receiving dialysis has historically been reported to be approximately 10–20-fold higher than that in the general population, and cardiovascular disease remains a major cause of death in this population [3]. Recent data from the United States Renal Data System (USRDS) further demonstrate the persistently high mortality associated with dialysis. Among Medicare fee-for-service beneficiaries aged ≥66 years in 2023, unadjusted mortality rates for dialysis patients ranged from approximately 244 to 370 deaths per 1,000 person-years depending on age and socioeconomic status, substantially exceeding those observed in kidney transplant recipients [4].

ESKD also imposes a substantial economic burden on healthcare systems. According to USRDS data, Medicare expenditures for dialysis treatment in the United States reached approximately US $45.3 billion in 2022 [5]. In the same year, annual per-person Medicare fee-for-service expenditures for patients with ESKD were approximately US $67,320 for Medicare-only beneficiaries and US $98,985 for Medicare–Medicaid dual-eligible beneficiaries [5]. These considerable healthcare expenditures underscore the need for KRT technologies that improve not only treatment efficiency but also patient mobility, quality of life, and independence from healthcare infrastructure.

The principal KRT options currently available to patients with ESKD are kidney transplantation, hemodialysis (HD), and peritoneal dialysis (PD). Kidney transplantation generally provides superior long-term survival, quality of life, and cost-effectiveness compared with maintenance dialysis; however, its availability remains constrained by donor shortages, transplant eligibility, and infrastructure requirements [6]. Consequently, a large proportion of patients with ESKD remain dependent on long-term HD or PD.

HD is the most widely used dialysis modality and removes urea, creatinine, electrolytes, and excess fluid from the blood across a semipermeable dialysis membrane. Conventional maintenance HD is typically performed three times per week for approximately 4 hours per session [7]. Because solutes and fluid accumulate during the interdialytic interval, substantial amounts must be removed over a relatively short treatment period. The resulting rapid changes in intravascular volume, electrolyte concentrations, and acid–base balance can contribute to intradialytic hypotension, myocardial stress, and recurrent cardiovascular instability. Thus, despite its high solute-clearance capacity, intermittent HD does not fully reproduce the continuous and gradual filtration provided by native kidneys.

By contrast, PD uses the patient’s own peritoneal membrane as a semipermeable barrier for solute and water transport between the peritoneal microcirculation and dialysate. Low-molecular-weight solutes, including urea, creatinine, potassium, and phosphate, are transported primarily by diffusion over several hours, whereas osmotic gradients generated by glucose-containing dialysate drive ultrafiltration of excess water. Because PD provides relatively slow and prolonged treatment, it more closely approximates continuous kidney filtration and generally produces smaller acute hemodynamic perturbations than intermittent HD. PD is also associated with preservation of residual kidney function and provides greater flexibility for home-based treatment, offering an important therapeutic option for patients seeking to maintain social and occupational activities.

Continuous ambulatory peritoneal dialysis (CAPD), one of the most widely established PD modalities, typically involves instillation of approximately 2 L of fresh dialysate into the peritoneal cavity, followed by a dwell period of several hours and subsequent drainage; this process is commonly repeated approximately four times per day. However, each exchange requires the patient to connect, drain, infuse, and dispose of dialysate bags, imposing substantial time and physical burdens. Repeated connections can also increase opportunities for touch contamination and PD-associated peritonitis [8]. Furthermore, chronic exposure to glucose-based PD solutions has been associated with systemic glucose absorption, weight gain, dyslipidemia, insulin resistance, and progressive alterations in peritoneal membrane structure and function . Automated peritoneal dialysis (APD) reduces some of the manual burden by using a cycler to perform multiple exchanges, usually during sleep, but still requires substantial volumes of fresh dialysate and storage of used fluid and restricts mobility during treatment.

Accordingly, a wearable continuous dialysate-regeneration system that minimizes repeated dialysate exchange while retaining the physiological advantages of PD represents an important goal for next-generation KRT. An ideal system would continuously withdraw used dialysate from the peritoneal cavity, selectively remove accumulated uremic solutes, and return regenerated dialysate to the patient, thereby enabling prolonged treatment using only a small dialysate volume. Such a strategy could reduce the frequency of dialysate-bag exchanges and improve patient mobility while maintaining slow and continuous solute and fluid removal, thereby limiting large temporal fluctuations in circulating uremic toxins and body fluid volume [9].

A critical requirement for realizing such a wearable system is a compact regeneration technology capable of continuously removing a broad range of uremic solutes from used dialysate. Biochemical adsorption using activated carbon or other sorbents, enzyme-mediated degradation, and membrane-based physical filtration have been investigated for dialysate regeneration. However, sorbent saturation, membrane fouling or clogging, cartridge replacement, and limited selectivity can constrain long-term continuous operation and device miniaturization [9]. In particular, a wearable platform must simultaneously process small ionic species and electrically neutral uremic molecules under strict constraints on filter volume and energy consumption. Electrochemical separation technologies, including electrodialysis (ED) [10,11], electrodeionization (EDI) [12], and capacitive deionization (CDI) [13,14], have therefore attracted considerable interest for compact and distributed water purification because of their modularity and potentially high energy efficiency [15]. Nevertheless, these technologies are predominantly optimized for ionic separation, making simultaneous treatment of solutes spanning markedly different sizes and electrochemical properties challenging.

To address these limitations, we previously developed an electrokinetic dialysis approach based on ion concentration polarization (ICP) [16]. ICP is an electrokinetic phenomenon generated by the permselectivity of nanoporous ion-selective membranes, producing local ion-enrichment and ion-depletion zones adjacent to the membrane [17–21]. The resulting strong localized electric field and ion-depleted region can be exploited to manipulate and separate charged species without relying solely on size-selective physical filtration [22,23]. In our previous work, this principle enabled continuous processing of cationic, anionic, and neutral species, and the achievable flow rate was increased from the microfluidic regime of approximately 0.1 μL/min to approximately 1 mL/min [16]. In this work, we present a conceptual backpack-type wearable platform for continuous regeneration of peritoneal dialysate (Fig. 1a), illustrating how the individual components evaluated in this study could be integrated into a compact recirculating system. The system integrates a nano-ring dialyzer, dialysate circulation pumps, sensing modules, supplementary and waste reservoirs, and a battery into a single wearable platform, enabling used dialysate withdrawn from the peritoneal cavity to be regenerated and continuously recirculated. A modular cascaded nano-ring dialyzer was developed to increase treatment capacity through parallelization (Fig. 1b), and two regeneration channels were designed to share a single waste channel to increase device integration density (Fig. 1c, Supplementary Fig. 1). The nano-ring mesh, which constitutes the core purification element, expands the ion-depletion zone and the spatial extent of electrokinetic transport, thereby enhancing the migration of charged uremic solutes. For electrically neutral urea, electroosmotic mixing within the microchannels of the nano-ring structure enhances mass transport toward the electrode surface, facilitating electrochemical removal (Fig. 1d,e).

**Fig. 1.**
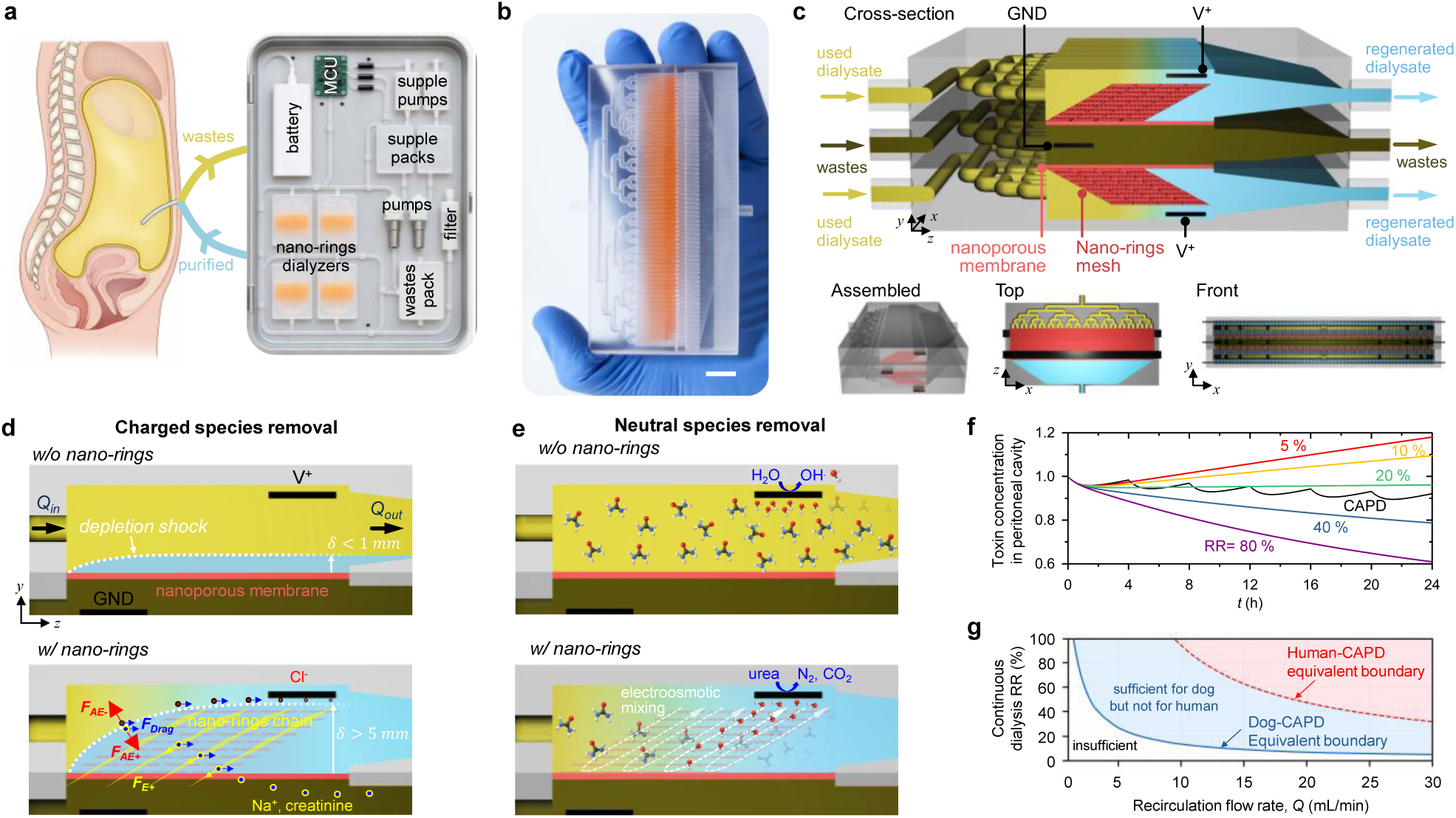
Concept and operating principle of the wearable continuous peritoneal dialysate-regeneration system. **a**, Conceptual illustration of a backpack-type wearable platform integrating a nano-ring dialyzer, dialysate circulation pumps, sensing modules, supplementary and waste reservoirs, and a battery for continuous regeneration and recirculation of peritoneal dialysate. **b,** Modular nano-ring dialyzer architecture designed to increase treatment capacity through parallelization. Inset scale bar, 1 cm. **c,** Integrated fluidic configuration in which two regeneration channels share a common waste channel to improve device integration density. **d,e,** Schematic illustration of the nano-ring mesh and its electrokinetic purification mechanism. The nano-ring structure expands the ion-depletion zone and enhances electrokinetic transport of charged uremic solutes, while electroosmotic mixing within the nano-ring microchannels promotes mass transport of electrically neutral urea toward the electrode surface for electrochemical removal. **f**, Biomimetic transport model incorporating continuous generation of urea and creatinine, transperitoneal solute transport, dialysate recirculation, and dialyzer removal efficiency to compare continuous regenerative peritoneal dialysis with conventional continuous ambulatory peritoneal dialysis (CAPD) in a renal-failure canine model. **g,** CAPD-equivalent operating map showing the combinations of dialysate recirculation flow rate and single-pass toxin removal efficiency required to achieve toxin-removal performance comparable to canine and human CAPD. At a dialysate flow rate of 10 mL/min, a removal efficiency above approximately 20 % was predicted to achieve canine CAPD-equivalent performance, whereas higher performance would be required to reach the corresponding human-scale CAPD condition.

We further developed a biomimetic transport model incorporating continuous generation of urea and creatinine, transperitoneal solute transport, dialysate recirculation, and dialyzer removal efficiency to quantitatively compare the systemic toxin-removal performance of continuous regenerative PD using the nano-ring dialyzer with conventional CAPD in a renal-failure canine model (Fig. 1f). At a dialysate flow rate of 10 mL/min, the model predicted that a single-pass toxin removal efficiency of >20 % would provide performance comparable to CAPD. The measured urea and creatinine removal capabilities of the nano-ring dialyzer were predicted to satisfy the CAPD-equivalent operating range for the canine model, whereas further improvement would be required to achieve the corresponding human-scale CAPD performance (Fig. 1g). Guided by these predictions, we fabricated a modular nano-ring dialyzer capable of operating at 10 mL/min and targeted a toxin removal efficiency exceeding 20 % for subsequent validation of continuous regenerative PD in a canine model.

## RESULTS

### Simulation-guided electrokinetic optimization of nano-ring dialyzer geometry

First, the effects of the structural parameters of the nano-rings mesh on the expansion of the ion-depletion zone and on solute removal performance were investigated using numerical simulations (Fig. 2a). Used dialysate introduced into the device sequentially passes through multiple cascaded Nano-ring structures. In the front view, nanoporous resin is coated around the openings of the mesh lattice, forming a ring-shaped pattern. Simulation details were shown in Supplementary Fig. 2.

**Fig. 2.**
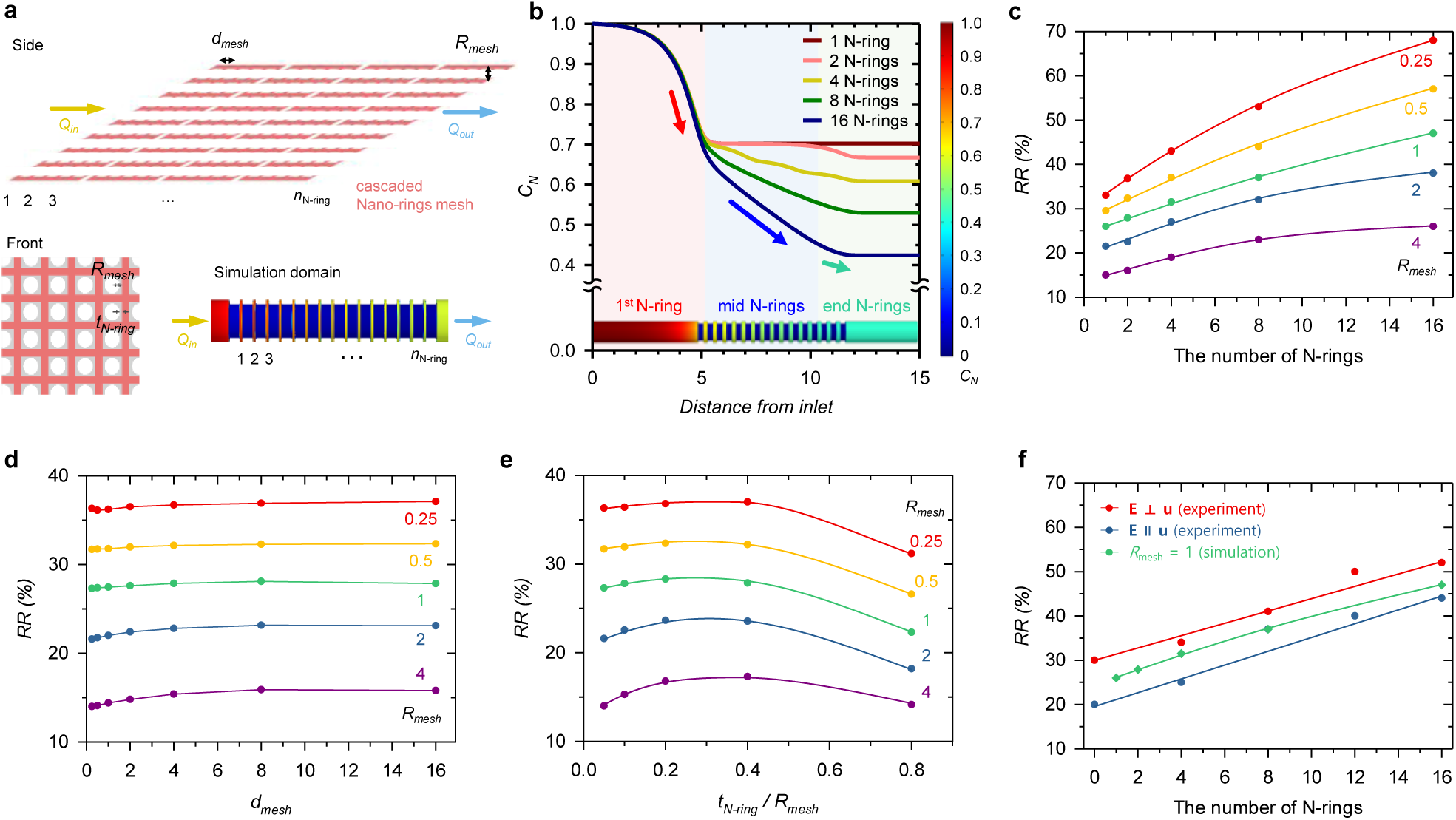
Structural optimization and electrokinetic deionization characteristics of the nano-rings mesh. **a,** Schematic illustration of the nano-ring mesh geometry and structural parameters used for numerical analysis. Used dialysate sequentially passes through multiple cascaded nano-ring-coated meshes, in which nanoporous resin forms ring-shaped patterns around the mesh openings. **b,** Simulated concentration distributions showing the progressive downstream expansion of the ion-depletion region with increasing numbers of cascaded nano-rings (N-rings). **c,** Removal ratio (*RR*) as a function of the number of nano-rings under different mesh-dimension conditions, showing enhanced cumulative removal with increasing nano-ring number and smaller mesh dimensions. **d,** Dependence of RR on mesh dimension and inter-mesh spacing, demonstrating higher removal for smaller mesh dimensions and only a minor influence of spacing between adjacent nano-ring-coated meshes. **e,** Effect of nano-ring coating thickness on *RR*, revealing a non-monotonic dependence with an intermediate optimum thickness. **f,** Experimental and simulated comparison of solute removal under different relative orientations between the flow direction and the applied electric field. Higher removal was obtained when the electric field was oriented perpendicular to the flow, consistent with enhanced transverse electrokinetic transport.

Comparison of the concentration distributions as a function of the number of nano-rings showed that a pronounced concentration decrease occurred near the first nano-ring, followed by a progressive downstream expansion of the ion-depletion region as additional nano-rings (N-rings) were introduced (Fig. 2b). With a single nano-ring, solute depletion remained largely confined to the region near the initial structure. By contrast, increasing the number of nano-rings caused the local depletion zones generated at individual rings to overlap and extend along the flow pass, resulting in a lower outlet concentration. In particular, as *N* increased, continued concentration reduction was observed around the intermediate and downstream nano-rings, indicating that a cascaded nano-ring configuration can extend a locally generated ion-depletion effect over a substantially larger portion of the flow channel. The effect of nano-ring number on removal performance was further quantified using the removal ratio (*RR*). *RR* increased continuously with increasing *N* under all tested *R_mes_*_ℎ_ conditions (Fig. 2c). At a given *N*, smaller *R_mes_*_ℎ_ consistently produced higher *RR* values, and this trend was maintained as the number of nano-rings increased. These results indicate that the local ion-depletion effects generated at individual rings accumulate along the cascaded structure, thereby enhancing overall solute removal. At the same time, mesh dimension acts as a key structural parameter that modulates the magnitude of this cumulative effect.

The influence of mesh dimension *R_mes_*_ℎ_ was also pronounced. For the same nano-ring configuration, smaller *R_mes_*_ℎ_ values resulted in higher *RR*, whereas *RR* decreased as the mesh dimension increased (Fig. 2d). This behavior can be attributed to the larger fraction of the flow cross-section occupied by the electric-field-affected and ion-depleted regions in narrower mesh geometries. Under these conditions, a greater proportion of the transported solute is exposed to strong electrokinetic effects. In contrast, as *R_mes_*_ℎ_ increases, the depletion layer occupies a smaller relative fraction of the flow path, reducing the overall removal efficiency.

In contrast, the spacing *L* between two nano-ring-coated meshes showed little correlation with removal performance within the tested range. This suggests that, under the present operating conditions, the cumulative contribution of the individual nano-ring elements is more strongly governed by the number and geometry of the coated meshes than by the separation distance between adjacent elements.

Nano-ring coating thickness *t*was also identified as an important structural parameter. *RR* initially increased with increasing *t_N_*_−*ring*_, reached a maximum at an intermediate thickness, and subsequently decreased with further increases in coating thickness (Fig. 2e). This non-monotonic behavior suggests the existence of an optimal coating geometry. When the coating is too thin, the resulting ion-depletion zone is insufficiently developed, whereas beyond the optimum thickness, additional coating alters the local electric-field and ion-transport distributions without providing further improvement in removal performance. Thus, solute removal cannot be maximized simply by increasing the coating thickness; instead, an appropriate *t* must be selected to achieve the most favorable electrokinetic transport condition.

Finally, the effect of the relative orientation between the flow direction and the applied electric field was evaluated experimentally and numerically (Fig. 2f). Higher removal was observed when the flow direction and electric field were oriented perpendicular to each other than when they were aligned in parallel, and the same trend was reproduced in the simulations. This behavior is attributed to enhanced transverse transport of solutes across the main flow direction when the electric field is applied orthogonally, thereby increasing solute interaction with the nano-ring structure and the ion-depletion region. By contrast, under parallel alignment, electrically driven transport is directed primarily along the bulk flow, limiting transverse redistribution. These results demonstrate that nano-ring-mediated removal is governed not only by flow rate and structural geometry but also by the directional coupling between hydrodynamic transport and the applied electric field.

Overall, these results establish the nano-ring mesh as a geometrically tunable electrokinetic purification architecture in which multiple local ion-depletion zones can be cascaded to extend solute depletion over the entire flow path. Increasing the number of nano-rings and reducing the mesh dimension enhanced cumulative removal, whereas coating thickness exhibited an optimum rather than a monotonic dependence, and inter-mesh spacing had only a minor effect. Moreover, the superior performance obtained under orthogonal flow–electric-field alignment highlights the importance of transverse electrokinetic transport. Together, these findings identify nano-ring number, mesh dimension, coating thickness, and field–flow orientation as the principal design parameters governing removal efficiency and provide a structural basis for optimizing the nano-ring dialyzer for continuous dialysate regeneration.

### Closed-loop continuous peritoneal dialysis using a nano-ring dialyzer

In this study, we established a closed-loop continuous peritoneal dialysis platform based on a nano-ring dialyzer and evaluated its dialysate regeneration and uremic toxin removal performance using both *in vitro* (Supplementary Fig. 3) and canine *in vivo* models (Supplementary Fig. 4).

We first assessed the purification characteristics of the cascaded nano-ring dialyzer using used peritoneal dialysate collected from patients undergoing peritoneal dialysis. As shown in Fig. 3a, the conductance difference between the regenerated and waste streams progressively increased with increasing applied voltage, indicating enhanced separation of ionic species under stronger electric fields. In contrast, only a marginal conductance difference was observed between the two outlet streams in a control device without nano-rings. These results demonstrate that the nano-ring architecture effectively enhances nanoelectrokinetics-mediated ionic separation and dialysate regeneration [24].

**Fig. 3.**
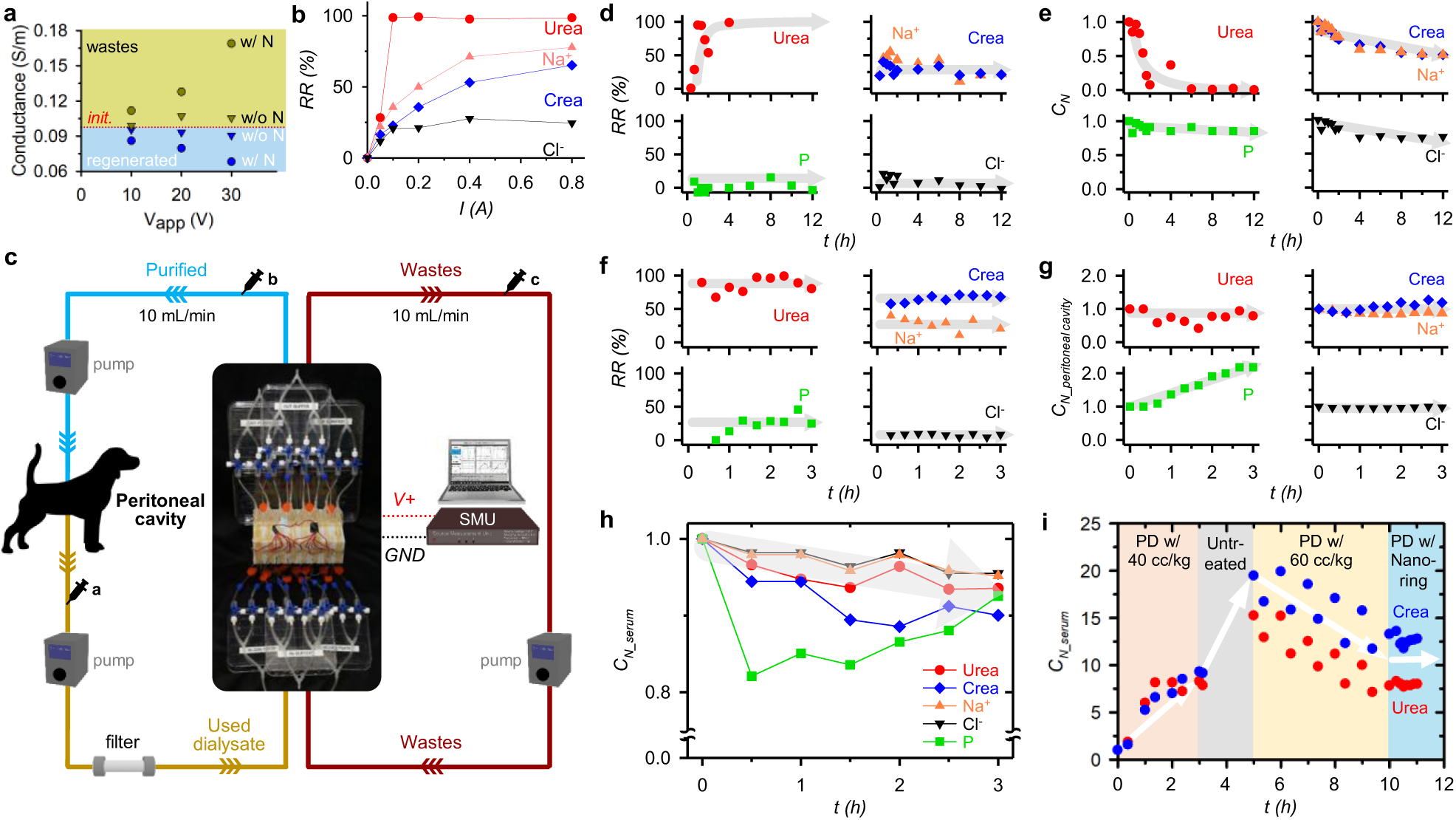
*In vitro* and *in vivo* validation of closed-loop dialysate regeneration using the nano-ring dialyzer. **a,** Conductance difference between regenerated and waste streams as a function of applied voltage using used peritoneal dialysate, comparing nano-ring and control devices. **b,** Single-pass removal efficiencies of urea and creatinine measured at a flow rate of 1.33 mL/min under increasing applied current. Removal increased with current and approached a plateau above approximately 0.4 A; 0.2 A was selected for subsequent closed-loop experiments. **c,** Parallel integration of eight nano-ring dialyzer modules to achieve a total processing capacity of approximately 10 mL/min, together with the associated pumping, filtration, sensing, sampling, and power components used for closed-loop operation. **d,e,** *In vitro* closed-loop recirculation experiments over 12 h showing single-pass removal efficiencies and temporal changes in solute concentrations within the peritoneal-mimicking reservoir. Urea was rapidly depleted, whereas creatinine and major electrolytes showed more gradual concentration changes. **f,** Single-pass removal efficiencies measured during *in vivo* continuous peritoneal dialysis in a canine renal-failure model, including urea, creatinine, Na⁺, Cl⁻, and phosphorus. **g,** Temporal changes in solute concentrations in peritoneal dialysate during continuous *in vivo* regeneration, reflecting simultaneous toxin removal by the dialyzer and solute transport from blood into the peritoneal cavity. **h,** Temporal changes in serum urea, creatinine, electrolytes, and phosphorus during continuous treatment, showing an overall reduction in circulating uremic toxin levels during operation. **i,** Long-term comparison of conventional peritoneal dialysis and nano-ring dialyzer-based continuous-flow PD in a bilateral-nephrectomy canine model over 12 days, demonstrating stabilization of serum urea and creatinine during continuous dialysate regeneration.

The uremic toxin removal performance of the nano-ring dialyzer was then quantitatively evaluated (Fig. 3b). Used and fresh dialysates were introduced into the anode-side and cathode-side channels, respectively, at a flow rate of 1.33 mL/min. Applied current was varied from 0.05 to 0.8 A, corresponding to steady-state voltages of approximately 5.8–25.3 V. Regenerated dialysate was sampled every 20 min over a total operating period of 60 min. Near-complete urea removal was observed at currents of 0.1 A or higher, whereas creatinine removal exceeded 30 % at currents above 0.2 A. The removal efficiencies of both solutes increased with applied current but tended to plateau above approximately 0.4 A. Based on the combined consideration of toxin removal efficiency and power consumption, 0.2 A was selected as the operating condition for subsequent closed-loop experiments, providing more than 30 % toxin removal per circulation cycle.

To increase treatment throughput, eight nano-ring dialyzer modules were integrated in parallel, yielding a total processing capacity of approximately 10 mL/min (Fig. 3c). The multi-module system was connected to peristaltic pumps controlling the regenerated dialysate, recirculating waste stream, and used dialysate, and the device was powered using a source measurement unit. A silk fibroin filter was additionally placed upstream of the nano-ring dialyzer to remove biological debris and enhance creatinine removal. Major dialysate constituents were monitored in real time using an integrated sensing unit, while dialysate and blood samples were periodically collected from designated sampling nodes and from the canine model for renal panel analysis.

Prior to *in vivo* testing, an *in vitro* closed-loop experiment was performed to mimic continuous peritoneal dialysis. A reservoir representing the peritoneal cavity was connected to the multi-module nano-ring dialyzer in a closed circuit, and the removal characteristics of urea, creatinine, and major electrolytes were evaluated over 12 hours. The single-pass removal ratio for each solute was calculated as *RR* = (1 − *C_b_*/*C_a_*) × 100 %, where *C_a_* and *C_b_* denote the solute concentrations measured upstream and downstream of the dialyzer, respectively. Urea exhibited a very high single-pass removal efficiency from the beginning of the experiment and was nearly completely removed within approximately 4 hours. The average creatinine removal ratio was approximately 27.8 %, close to the design target of 30 %. The removal efficiencies of major electrolytes varied among species but were generally lower than those of urea and creatinine. This selective removal behavior suggests that the nano-ring dialyzer can effectively remove major uremic toxins while limiting excessive loss of essential electrolytes.

Analysis of solute concentrations within the peritoneal-mimicking reservoir showed that urea concentration decreased sharply during the first 2 hours, approaching near-complete removal. This behavior was attributed to the sustained high urea removal efficiency of the multi-module nano-ring dialyzer during repeated circulation. Creatinine and electrolyte concentrations decreased more gradually and progressively declined over 12 hours of recirculating purification (Fig. 3d,e). The *in vitro* closed-loop experiments were repeated more than ten times to confirm the reproducibility of the system.

Based on these results, an *in vivo* closed-loop continuous PD experiment was conducted in a canine model of chronic renal failure. Following placement of an intraperitoneal catheter, spent dialysate was continuously withdrawn, regenerated through the nano-ring dialyzer, and recirculated into the peritoneal cavity. Fig. 3f shows the single-pass removal ratios measured under *in vivo* conditions. Urea removal remained high, with an average efficiency of 86.4 %, while creatinine removal averaged 66.1%. The average removal ratios of Na⁺ and Cl⁻ were 28.3% and 7.7%, respectively. Notably, creatinine removal was substantially higher than that observed *in vitro*, which was likely aided by the upstream silk fibroin filter. Phosphorus removal was also enhanced relative to the *in vitro* condition, reaching an average of approximately 20.3 %.

Fig. 3g shows the temporal changes in solute concentrations within the peritoneal dialysate during continuous treatment. Urea concentration gradually decreased, whereas creatinine and phosphorus showed modest increases over time. In contrast, Na⁺ and Cl⁻ concentrations remained relatively stable. These trends indicate that uremic toxins were continuously removed by the nano-ring dialyzer while newly generated solutes were simultaneously transported from the bloodstream into the peritoneal dialysate through diffusion and convection across the peritoneum. The relatively stable electrolyte concentrations further suggest that solute exchange and electrolyte balance were maintained during continuous dialysate regeneration.

Changes in major serum markers were simultaneously monitored in the canine model (Fig. 3h). As regenerated dialysate was continuously reintroduced into the peritoneal cavity, serum urea and creatinine concentrations gradually decreased, while Na⁺ and Cl⁻ also showed mild downward trends. Serum phosphorus generally decreased, although transient fluctuations were observed at several time points, potentially reflecting contributions from cell injury or hemolysis during the experiment. Overall, the major circulating uremic toxin levels decreased by approximately 10 % during 3 hours of continuous operation. These results demonstrate that the multi-module nano-ring dialyzer can continuously extract uremic solutes from the peritoneal cavity and reduce their systemic concentrations under *in vivo* conditions.

To further evaluate the potential of nano-ring-based continuous PD for prolonged treatment, a 12-day *in vivo* study was conducted in a canine model following bilateral nephrectomy (Fig. 3i). During the first 3 days after nephrectomy, conventional PD was performed using 40 cc/kg of dialysate, during which serum urea and creatinine concentrations continued to increase. Both toxin levels rose more rapidly during the subsequent 2-day treatment-free period. Conventional PD using 60 cc/kg of dialysate was then performed for 5 days, resulting in a decrease in serum urea and creatinine concentrations. The same animal was subsequently treated using nano-ring dialyzer-based continuous-flow PD. Under this condition, serum urea and creatinine concentrations remained relatively stable without further rapid accumulation and were maintained near the preceding baseline range. These results suggest that continuous regeneration and recirculation of a limited dialysate volume using the nano-ring dialyzer may provide uremic toxin control comparable to that achieved using conventional intermittent PD with repeated administration of 40–60 cc/kg of dialysate.

Collectively, these results demonstrate that the nano-ring dialyzer provides high single-pass uremic toxin removal, scalable treatment throughput through parallel module integration, and sustained dialysate regeneration within a closed-loop system. Importantly, effective control of circulating uremic toxin levels was demonstrated in canine renal-failure models, supporting the potential of an integrated system combining the nano-ring dialyzer with compact pumps, filters, sensors, and power modules as a platform for future portable or wearable continuous peritoneal dialysis.

### Improved biocompatibility for prolonged continuous peritoneal dialysis

Whereas the preceding section focused on the uremic toxin removal capability of the nano-ring dialyzer and its feasibility for continuous peritoneal dialysis, this section addresses additional safety requirements for prolonged *in vivo* use, including pH stabilization, mitigation of oxidative byproducts, suppression of biological contamination, and improvement of overall biocompatibility.

Because the conventional nano-ring dialyzer relies on an anion-selective nanoporous membrane, preferential transport and removal of cationic species can occur, potentially disturbing ionic balance and shifting the pH of the regenerated dialysate. To mitigate this effect, the conventional wire-type electrodes were replaced with plate-type electrodes, increasing the effective electrode–fluid contact area by more than tenfold (Fig. 4a). The enlarged electrode area was designed to generate a more uniform electric-field distribution across the nanomembrane and thereby reduce device-to-device variation in local field strength, current distribution, and flow behavior.

**Fig. 4.**
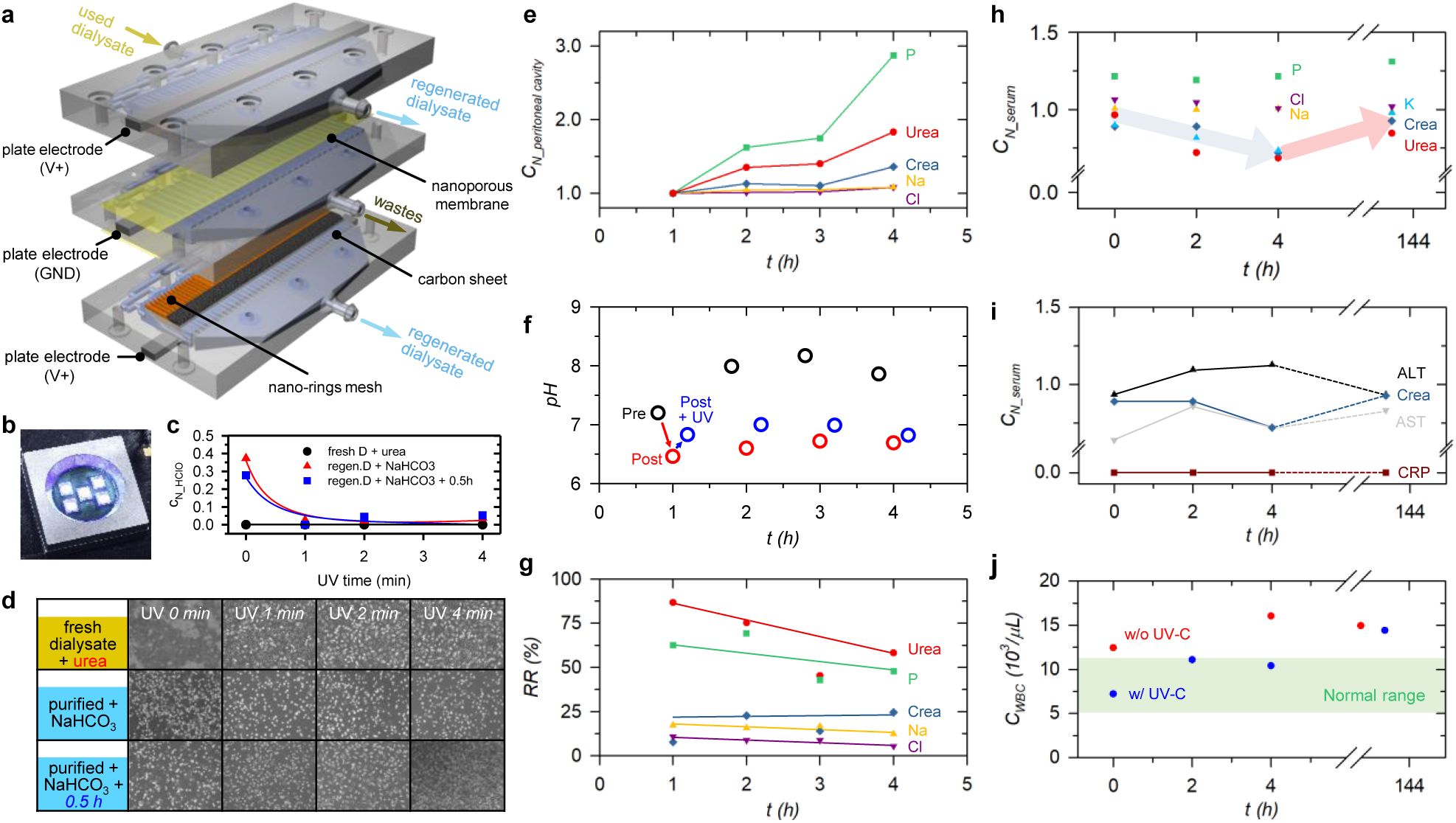
Improved biocompatibility and prolonged *in vivo* operation of the nano-ring-based continuous peritoneal dialysis system. **a,** Design modifications introduced to improve chemical and fluidic stability during prolonged operation, including replacement of wire electrodes with plate-type electrodes and incorporation of activated carbon adjacent to the nano-ring mesh and nanoporous membrane. **b,** Integration of UV-C treatment downstream of the nano-ring dialyzer for attenuation of reactive chemical byproducts. **c,** Reduction of oxidative species in regenerated dialysate following UV-C exposure, including HClO-related compounds. **d,** Cell viability after exposure to regenerated dialysate under different treatment conditions, showing improved biocompatibility following sequential pH neutralization, equilibration, and UV-C treatment. **e,** Temporal changes in solute concentrations in peritoneal dialysate during 4 h of continuous closed-loop PD in an unanesthetized canine model, reflecting ongoing blood-to-dialysate transport during recirculation. **f,** pH changes before and after the dialyzer and following UV-C treatment with NaHCO₃-based buffering. **g,** Single-pass removal ratios of BUN, creatinine, inorganic phosphorus, Na⁺, and Cl⁻ during continuous *in vivo* treatment. **h,** Temporal changes in serum solute concentrations during treatment and at 144 h after treatment, showing decreases in major circulating uremic toxins. **i,** Changes in serum ALT, AST, creatinine, and CRP during and after continuous PD, indicating the absence of sustained hepatic injury or persistent systemic inflammatory activation. **j,** White blood cell concentrations in peritoneal dialysate with and without UV-C treatment, showing reduced short-term inflammatory response during UV-C-assisted operation. Together, these results demonstrate that chemical conditioning, UV-C treatment, and system-level modifications improved the biocompatibility of regenerated dialysate while maintaining effective toxin removal during prolonged continuous PD.

In addition, activated carbon sheets were integrated adjacent to the Nanoporous resin-coated mesh and nanoporous membrane. While the nanoporous membrane structures primarily mediated cation exchange and transport, the activated carbon layer was introduced to adsorb anionic oxidative species and other reactive byproducts generated during purification, thereby helping to restore the ionic and chemical balance of the regenerated dialysate. To further compensate for device-induced acidification, a sodium bicarbonate (NaHCO₃)-based pretreatment step was incorporated to maintain the pH of regenerated dialysate within a range suitable for prolonged biological application.

A UV-C treatment module was subsequently introduced to further reduce potentially harmful oxidative byproducts (Fig. 4b). When regenerated dialysate collected after passage through the nano-ring dialyzer was exposed to UV-C irradiation, the levels of oxidative species, including hypochlorous acid (HClO)-related compounds, were markedly reduced after more than 1 min of treatment (Fig. 4c). These results indicate that UV-C exposure can attenuate residual reactive byproducts, potentially through photolytic decomposition or conversion to less reactive chemical species.

The cytotoxicity of regenerated dialysate was then evaluated by assessing cell viability 24 hours after exposure under different treatment conditions (Fig. 4d). Cells exposed to fresh dialysate exhibited normal growth, whereas untreated regenerated dialysate showed pronounced cytotoxicity. Simple pH neutralization alone was insufficient to restore cell viability, regardless of UV exposure time. In contrast, when neutralized regenerated dialysate was allowed to equilibrate for approximately 30 min before UV-C treatment, substantial recovery of cell viability was observed after 4 min of irradiation. This improvement is likely attributable to the time-dependent decay or transformation of transient reactive chemical species present in freshly regenerated dialysate, followed by further attenuation of residual oxidative byproducts during UV-C exposure. Immediately after neutralization, acid–base reactions and secondary reactions involving reactive species may remain incomplete. A short equilibration period may therefore allow unstable intermediates to dissipate or convert into more stable species, after which UV-C treatment can further reduce the remaining reactive components. These results suggest that a sequential neutralization–equilibration–UV-C treatment strategy is more effective than pH correction alone for improving the biocompatibility of regenerated dialysate.

Based on these findings, an improved closed-loop continuous PD system was assembled as shown in Supplementary Fig. 5 and evaluated in a canine *in vivo* model. In this experiment, the silk-fibroin auxiliary filter used in the previous section was omitted to assess the intrinsic performance of the nano-ring dialyzer. In addition, experiments were conducted in unanesthetized animals to more closely approximate the intended conditions for a portable continuous PD system.

Continuous peritoneal dialysis was performed for 4 hours. During treatment, the concentrations of uremic toxins and selected ions in dialysate collected from the peritoneal cavity gradually increased over time (Fig. 4e). This behavior indicates that, while regenerated dialysate was continuously reintroduced into the peritoneal cavity, urea, creatinine, and other solutes were continuously transferred from the bloodstream into the dialysate across the peritoneum by diffusion and convection. Thus, repeated recirculation of purified dialysate maintained the concentration gradient required for ongoing blood-to-dialysate solute transport.

To prevent excessive acidification of the regenerated dialysate, 4.5 g/L NaHCO₃ solution was introduced before the dialyzer. This treatment maintained the post-dialyzer pH above 6, while an additional modest increase in pH was observed following UV-C treatment (Fig. 4f). These results suggest that the combination of bicarbonate buffering and downstream UV-C treatment contributes to maintaining acid–base stability during prolonged recirculation.

Single-pass removal ratios were calculated from the concentration differences measured before and after the dialyzer (Fig. 4g). BUN, creatinine, inorganic phosphorus (IP), Na⁺, and Cl⁻ exhibited removal ratios of approximately 50–80 %, 10–25 %, 40–70 %, 10–20 %, and 5–10 %, respectively. The removal efficiencies of urea and phosphorus gradually declined over time, whereas creatinine removal showed a modest increase. In contrast, Na⁺ and Cl⁻ removal remained relatively stable throughout the experiment. These findings demonstrate that the nano-ring dialyzer can sustain removal of major uremic solutes even in the absence of an auxiliary filter, while limiting excessive electrolyte loss.

Fig. 4h shows changes in serum composition during continuous dialysis as well as serum concentrations measured 144 hours after the experiment in the surviving canine. During the 4-h continuous PD treatment, the concentrations of major uremic toxins in serum generally decreased. In particular, urea and creatinine showed clear downward trends, directly demonstrating that continuous recirculation of regenerated dialysate can reduce circulating toxin levels. Although phosphorus exhibited substantial removal across the dialyzer, its serum concentration remained relatively stable, possibly because of ongoing redistribution between intracellular or tissue compartments and the bloodstream. By 144 hours after treatment, most serum analytes had returned to levels close to their pre-treatment values. To further assess biological safety, alanine aminotransferase (ALT), aspartate aminotransferase (AST), creatinine, and C-reactive protein (CRP) were monitored (Fig. 4i). ALT and AST are commonly used biochemical markers of hepatocellular injury and liver-associated tissue damage, respectively, whereas creatinine reflects renal clearance and systemic accumulation of nitrogenous waste products. CRP is a widely used acute-phase marker of systemic inflammation. During continuous PD, serum creatinine decreased modestly and returned to near-baseline levels at 144 hours. AST and CRP remained close to their initial values throughout the observation period. ALT transiently increased during treatment but returned to its initial level by 144 hours after completion of the experiment. The transient nature of this increase, together with subsequent recovery, suggests that the 4-h continuous treatment did not induce sustained hepatic injury or a persistent systemic inflammatory response. Finally, the potential for peritoneal inflammatory response was evaluated by monitoring white blood cell (WBC) concentrations in the peritoneal dialysate (Fig. 4j). In the absence of UV-C treatment, WBC levels increased after approximately 4 hours of treatment. In contrast, under UV-C-treated conditions, only a modest increase was observed, and WBC levels remained within the normal range during the 4-h treatment period. This finding suggests that UV-C treatment may help reduce short-term biological or inflammatory stimulation within the recirculating circuit. However, by 144 hours, WBC levels in the UV-C-treated group had increased to values comparable to those in the untreated condition, indicating that long-term suppression of peritoneal inflammation will likely require additional strategies, including improved circuit sterility, catheter management, and infection-control measures.

Collectively, the improved nano-ring dialysis system maintained uremic toxin removal while simultaneously enhancing pH stability, reducing reactive chemical byproducts, and limiting short-term inflammatory responses. The combined use of plate-type electrodes, activated carbon-assisted chemical conditioning, bicarbonate buffering, and UV-C treatment mitigated several chemical and biological instabilities that could otherwise limit prolonged *in vivo* application of the nano-ring dialyzer. Importantly, continuous closed-loop PD was successfully performed for 4 hours in unanesthetized canines without the use of the auxiliary silk-fibroin filter, while serum uremic toxin levels decreased and major hepatic and inflammatory biomarkers remained within recoverable ranges. These results support the potential of the nano-ring platform to evolve from a dialysate purification device into a regenerative peritoneal dialysis system suitable for prolonged *in vivo* operation.

Finally, all major components used in the *in vivo* experiments were integrated into a single portable bag-type configuration (Supplementary Fig. 6). The integrated structure measured 55 cm × 35 cm × 5 cm and had a total mass of 4.7 kg, including a 3-kg casing (Supplementary Fig. 6a). A total of ten functional modules were successfully incorporated into the system (Supplementary Fig. 6b), and the overall load was experimentally confirmed to be compatible with adult carrying capacity (Supplementary Fig. 6c). This integration demonstrates the feasibility of combining the nano-ring dialyzer, circulation pumps, chemical conditioning modules, UV-C treatment unit, sensing components, and battery within a single mobile platform. With further validation of long-term safety, purification efficiency, infection control, and user compatibility in future preclinical and clinical studies, this integrated system may provide a foundation for portable or wearable peritoneal dialysis based on continuous dialysate regeneration.

## DISCUSSION

The present study suggests that nano-ring-assisted ion concentration polarization can serve as a practical basis for continuous dialysate regeneration in peritoneal dialysis. The key advantage of the nano-ring architecture lies in its ability to extend localized ion-depletion phenomena along the flow pass, thereby converting a confined electrokinetic effect into a spatially distributed purification process. This behavior indicates that device performance is governed not only by the applied electrical input, but also by geometric factors that control local field distribution, depletion-zone development, and solute transport.

The non-monotonic dependence on nano-ring coating thickness further suggests that purification efficiency is determined by a balance between ion-selective transport and local electric-field configuration rather than by membrane area alone. Likewise, the higher performance obtained when the electric field was oriented transverse to the flow supports the importance of directional electrokinetic transport. These observations emphasize that nano-ring dialyzer design should be treated as a coupled optimization problem involving geometry, fluid flow, and electric-field orientation.

A key advantage of closed-loop continuous regeneration is that a limited volume of dialysate can be continuously purified and recirculated, thereby maintaining the concentration gradient between the blood and dialysate and sustaining uremic toxin transport across the peritoneum. Unlike conventional PD, which relies on repeated replacement of fresh dialysate, this approach provides a treatment mechanism that more closely mimics the continuous clearance function of the native kidney.

The *in vivo* observations also highlight the importance of considering the entire biological transport system rather than the dialyzer alone. Solute concentrations within the peritoneal cavity reflect the dynamic balance between membrane transport, dialysate recirculation, and device-mediated removal. Accordingly, the effectiveness of continuous PD should ultimately be evaluated in terms of whole-body toxin control, fluid balance, and electrolyte homeostasis rather than single-pass removal efficiency alone.

For prolonged application, chemical compatibility of regenerated dialysate remains a critical challenge. The observed need for pH correction and treatment of reactive byproducts indicates that electrokinetic purification can introduce secondary chemical changes that must be actively managed. The beneficial effect of combining buffering, equilibration, activated carbon, and UV-C treatment suggest that dialysate regeneration should be regarded as a multistep conditioning process rather than a single purification step. In particular, the persistence of cytotoxicity after pH neutralization alone implies that transient reactive species may contribute independently to biological compatibility.

The short-term stability of major biochemical and inflammatory markers is encouraging, but long-term safety cannot yet be inferred. Future studies should directly characterize reactive byproducts, membrane fouling, electrolyte drift, catheter-associated infection risk, and tissue-level responses during prolonged operation. Closed-loop sensing and automated feedback control will likely be essential for maintaining pH, electrolyte composition, and treatment intensity over extended periods. Another important aspect of the platform is its modular scalability. Parallel integration of nano-ring units provides a practical route to increasing throughput without fundamentally altering the electrokinetic behavior of each unit. This modularity may enable treatment capacity to be adjusted according to patient size, residual renal function, and required clearance. At the same time, reduction in power consumption, module volume, and auxiliary fluid requirements will be necessary for true wearable implementation.

Overall, the significance of this work lies not only in demonstrating toxin removal, but in establishing a system-level framework for regenerative peritoneal dialysis based on continuous dialysate reuse. The combination of geometry-controlled electrokinetic purification, modular scale-up, chemical conditioning, and closed-loop circulation provides a foundation for a lower-intensity and more continuous form of kidney replacement therapy. With further optimization of safety, automation, and energy efficiency, this approach could contribute to the development of portable or wearable artificial kidney systems that more closely approximate the continuous clearance function of the native kidney.

## METHODS

### Biomimetic modeling of CAPD and continuous regenerative peritoneal dialysis

A compartment-based biomimetic model was developed to quantitatively compare conventional continuous ambulatory peritoneal dialysis (CAPD) with continuous regenerative peritoneal dialysis. The model incorporated endogenous generation of urea and creatinine, transperitoneal diffusive and convective solute transport, glucose-driven osmotic ultrafiltration, dialysate recirculation, and dialyzer-mediated solute removal. Body and dialysate compartments were described by time-dependent mass-balance equations for urea and creatinine, while dialysate glucose concentration and volume were simultaneously updated to account for glucose absorption and osmotic water transport. For each solute *s*, transperitoneal diffusive transport was described as

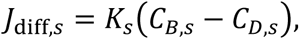

where *K_s_* is the effective peritoneal mass-transfer coefficient and *C_B_*_,*s*_ and *C_D_*_,*s*_ are the body and dialysate concentrations, respectively. Convective solute transport associated with positive ultrafiltration was expressed as

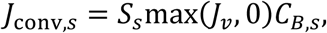

where *S_s_*is the sieving coefficient and *J_v_*is the net transperitoneal fluid flux. The body-compartment concentration was therefore governed by

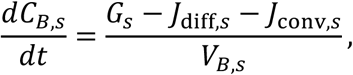

where *G_s_*represents endogenous solute generation and *V_B_*_,*s*_represents the apparent body distribution volume.

Glucose-dependent ultrafiltration was modeled from the osmotic pressure difference between dialysate and blood,

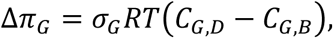

and the net fluid flux was calculated as

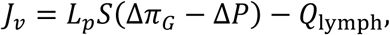

where *σ_G_*is the glucose reflection coefficient, *L_p_S* is the effective hydraulic permeability–surface area product, Δ*P* is the hydrostatic pressure difference, and *Q*_lymph_ is lymphatic fluid removal. Dialysate glucose was allowed to decrease through transperitoneal absorption during each treatment period.

For CAPD simulations, fresh dialysate was introduced at the beginning of each dwell, and the dialysate was completely replaced four times per day. Solute accumulated in the dialysate during each dwell and was removed at drainage. For the continuous regenerative system, the dialysate remained in continuous circulation through an external purifier. Purifier-mediated removal of urea and creatinine was modeled as

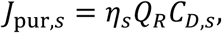

where *Q_R_* is the dialysate recirculation flow rate and *η_s_* is the single-pass dialyzer removal efficiency. Thus, increasing either recirculation rate or removal efficiency increased the effective regenerative clearance of the dialysate.

For the canine model, a body weight of 12 kg was used. Urea and creatinine distribution volumes were set to 0.60 *W*and 0.43 *W*, respectively, and the initial body concentrations were 15.0 mmol/L for urea and 0.80 mmol/L for creatinine. Endogenous generation rates were scaled by body weight from 70-kg reference values of 0.18 mmol/min for urea and 0.0060 mmol/min for creatinine. The peritoneal transport coefficients were set to 10 mL/min for urea and 8 mL/min for creatinine. The initial dialysate fill volume was set to 40 mL/kg. For comparison with a human-scale system, a 70-kg model with a dialysate fill volume of approximately 2.0 L was additionally simulated, using effective transport coefficients of 15 and 10 mL/min for urea and creatinine, respectively.

To compare overall toxin burden, urea and creatinine concentrations were normalized to their respective initial concentrations and combined into a composite toxin index,

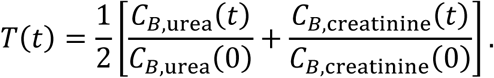

Time-dependent toxin profiles were calculated for CAPD and for continuous regenerative dialysis over a range of single-pass dialyzer removal efficiencies. In addition, a parametric sweep of recirculation flow rate and removal efficiency was performed. For each condition, the time-averaged urea and creatinine concentrations were normalized to the corresponding CAPD values and combined

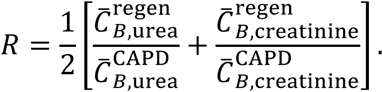

The contour defined by *R* = 1 was designated as the CAPD-equivalent boundary, representing combinations of dialysate recirculation rate and dialyzer removal efficiency that achieved the same time-averaged body toxin burden as conventional CAPD. Separate CAPD-equivalent contours were calculated for canine and human-scale models.

### Numerical simulation for nano-rings mesh

Ion transport through the nano-ring mesh was simulated in COMSOL Multiphysics using the General Form PDE interface. A two-dimensional longitudinal domain containing sequential cation-selective nano-ring membranes was considered. Ionic transport was described using a coupled Poisson– Nernst–Planck formulation, in which diffusion, electromigration, and convection were simultaneously considered. For a symmetric binary electrolyte, the dimensionless space-charge density was defined as

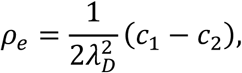

where *c*_1_and *c*_2_ are the dimensionless cation and anion concentrations and *λ_D_* is the dimensionless Debye length. The equilibrium ionic concentrations were initialized according to the Boltzmann distributions,

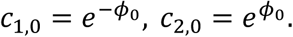

The fluid motion was modeled as incompressible creeping flow, with inertial terms neglected, by solving the Stokes equations,

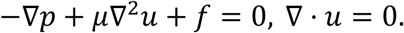

Linear finite elements were used for both velocity and pressure (*P*_1_ + *P*_1_), together with the stabilization implemented in COMSOL for equal-order interpolation. The resulting flow, electric-potential, and ionic-concentration fields were solved in a coupled manner to evaluate formation and downstream extension of the ion-depletion zone around the nano-ring structures.

### Building a nano-ring dialyzers

The design of the 3-D cascaded nano-ring dialyzers was created using a commercial 3-D drawing software (RhinoCeros 5.0). These devices consisted of distinct components including anodic compartments, a micro-mesh structure, a nanoporous membrane, and cathodic compartments, all illustrated in Fig. 1c and Fig. 4a. Here Nafion nanoporous membrane (with a thickness of 0.002 inches, sourced from Sigma Aldrich) and resin (20 w.t.% resin, Sigma Aldrich) was used in this work. The channel frames were manufactured using a Projet 2500+ 3D printer from 3Dsystems, USA, utilizing MJP RWT resin. Additionally, the mesh structure’s frame was 3-D printed using an M125 printer from MiiCraft, USA, employing photopolymer FC-2 from MiiCraft, USA.

### Apparatus for macro-fluidic experiment

Eight individual macro units were tightly assembled to form the final device, which exhibited a throughput of 10 mL/min. The process of unit integration involved utilizing non-toxic silicon adhesive as a bonding agent for each individual device. The interconnections between the device and the *in vitro* circuit were established through the utilization of Tigon tubes with an inner diameter of 3.2 millimeters, specifically sourced from ISMATEC.

To facilitate the necessary electrochemical processes, an external voltage was meticulously applied, a task accomplished through employment of a source measurement unit, namely the Lambda zup 36-6 from TDK. The controlled introduction of the analyte and buffer solutions (used dialysate from Seoul National University Hospital) was achieved through continuous injection, employing a consistent flow rate, into both the anodic and cathodic channels. This injection process was carried out by means of a custom-manufactured peristaltic pump obtained from Seoul National University Hospital, utilizing dialysate sourced from the same institution.

Subsequently, distinct streams of the injected solutions were extracted and isolated for analysis. Concentration profiles of specific ions including Na^+^ and Cl^−^, along with analytes like creatinine and urea, were quantitatively assessed using the Renal Panel method, facilitated by the HITACHI 7180 instrument.

### *In vitro* Closed-loop Continuous PD Using a Multimodule nano-ring Dialyzer

In the initial phase, a quantity of 2 L of previously employed dialysate was introduced into a chamber designed to mimic the peritoneal cavity. This chamber was used to replicate the volume of peritoneal fluid found within the body’s peritoneal cavity. In addition, we established a chamber that imitated the blood compartment, as well as an excess volume removal chamber. These chambers were utilized to replicate the transport of uremic toxins from the blood compartment into the peritoneal cavity through the peritoneum.

The transfer of used dialysate from the blood compartment mimic chamber to the peritoneal cavity mimic chamber occurred at a rate of 2 mL/min, while an excess volume of dialysate flowed from the peritoneal cavity mimic chamber into the excessive volume removal chamber at the same rate. This process ensured the maintenance of a constant total volume level within the peritoneal cavity mimic chamber. Meanwhile, the utilized dialysate present in the peritoneal cavity mimic chamber was directed into one side of a multimodule nano-ring dialyzer at a flow rate of 10 mL/min. From there, a purified dialysate was extracted, and subsequently reintroduced into the peritoneal cavity mimic chamber.

This cycle of dialysate circulation was perpetually repeated to effectively purify the previously used dialysate within the peritoneal cavity mimic chamber. Conversely, a buffer unit corresponding to the opposite side of the multimodule nano-ring dialyzer operated on an independent circuit. This unit contained an initial fill of fresh dialysate, which was continuously circulated at a rate of 10 mL/min. Samples at designated points labeled as “a” and “b” were collected at intervals of 20 minutes for the initial 2-hour period, and then at 2-hour intervals for the subsequent 10-hour duration.

### *In vivo* Canine Model Using Chronic Renal Failure Beagle Dogs

Male Beagle dogs in the adult stage (aged 14-16 months; weighing 8-11 kg) were employed to establish a model of chronic kidney disease through a 15/16 nephrectomy procedure. This involved ligating seven out of eight left renal arteries to induce partial infarction in the left kidney. Subsequently, the contralateral kidney was removed a week later, resulting in a remnant kidney comprising 1/16 of the original renal mass. After sixteen weeks from the second surgery, approximately 60% of the beagle subjects exhibited sustained elevated levels of serum creatinine (Scr > 2.0 mg/dl) and proteinuria (Protein to creatinine ratio > 1.5 g/gCr), effectively replicating the characteristics of chronic kidney disease.

Within this established model, two dogs underwent an additional procedure involving the insertion of peritoneal catheters. This operation was performed while the dogs were in a supine position. An incision was made on the anterior peritoneal wall on both sides to expose the peritoneal membrane. Dual-cuff Tenckhoff catheters were introduced bilaterally into the peritoneal cavity through these incision sites. A purse-string suture was applied to prevent any leakage of dialysate.

The experimental process commenced by introducing 600 mL (equivalent to 60 cc/kg) of fresh dialysate into the beagle dog’s peritoneal cavity. Through the peritoneum, an active exchange of substances occurred between the dialysate and the serum, facilitating the diffusion of body toxins from the serum into the peritoneal cavity. After a 4-hour period, the contaminated dialysate was removed from the peritoneal cavity using a peristaltic pump. The dialysate was initially passed through a filter, followed by a multimodule nano-ring dialyzer, for the purpose of purification. Notably, a silk-fibroin filter coated with Renamezin was employed. This filter type allowed *in vivo* debris to be filtered out of the dog’s peritoneal cavity, while Renamezin facilitated the adsorption of creatinine. The purified dialysate, having undergone this purification process, was then reinjected into the peritoneal cavity after passing through a sensor that monitored urea and electrolyte concentrations. To maintain a continuous PD process, a buffer unit was incorporated on the cathodic side, establishing an independent circuit that circulated additional used dialysate. Consequently, the purified dialysate was automatically reintroduced into the peritoneal cavity, enabling a continuous PD process. Notably, the setup involved three sampling points: one for the dog’s serum, the second for the exit of dialysate from the peritoneal cavity, and the third for the exit of the multimodule nano-ring dialyzer.

The experimental protocols have been approved by the Seoul National University Ethical Review Committee (under the approval code, PBT-19-039) with the relevant guidelines and regulations of the standards set by the Declaration of Helsinki.

## ACKNOWLEDGEMENTS

This work is mainly supported by Korean Health Technology RND project, Ministry of Health and Welfare Republic of Korea (HI14C0559020019) and partially supported by the Ministry of Science and ICT (#RS-2026-25607756 and #RS-2026-25502729, #NRF-2021R1C1C2012603). Also this work supported by BK21 four program. All *in vivo* experiments were approved by the Institutional Animal Care and Use Committee in Seoul National University Hospital (SNUH-IACUC) (IACUC No. 17-0221-S1AO) and animals were maintained in the facility accredited AAALAC International (#001169) in accordance with Guide for the Care and Use of Laboratory Animals 8th edition, NRC (2010).

## Competing Financial Interests

The authors declare no competing financial interests

## Author Contributions

W. Kim and K. Kim established the nano-ring dialysis system. W. Kim performed numerical simulations. H. Lee provided guidance on numerical simulations. K. Kim developed 3-D printed scaled-up device in macro-fluidic platform. S. Hong, M. Seo, D. Kim fabricated 3-D printed scaled-up device. W. Kim, K. Kim and S. Hong conducted *in vitro* and *in vivo* experiment. S. Lee, S. H. Yang, H. Lee, D. K. Kim contributed to make beagle model by performing operation of nephrectomy, PD catheter insertion and by conducting dialysis care. S. Lee, H. Lee, Y. S. Kim gave clinical analysis and guidance to overall *in vitro* and *in vivo* experiments. J. C. Lee and H. C. Kim designed experiment methods and conducted result interpretation, and D. A. Shin, K. J. Lee, W. S. Cho fabricated pump and circuit for experiments and conducted *in vitro* and *in vivo* experiments and data analysis. J. Lee and K. Kim conducted the urea sensor experiment for flowing sample. L. P. Lee provided strategic guidance on the overall direction of the study. G. Y. Sung and S. J. Kim supervised the project.

## Notes

### Competing Interest Statement

The authors have declared no competing interest.

