## Supplementary material for "Cascaded Nano-ring Dialyzer for Continuous Regenerative Peritoneal Dialysis": Supplemenatary Fig

Prof. Luke P. Lee, Prof. Yon Su Kim, Prof. Jung Chan Lee, Prof. Gun Yong Sung and Prof. Sung Jae Kim.

E-mail:

(Luke P. Lee)

(Yon Su Kim)

(Jung Chan Lee)

(Gun Yong Sung)

(Sung Jae Kim)

### **Supplementary Fig. 1.** Modular cascaded nano-ring dialyzer

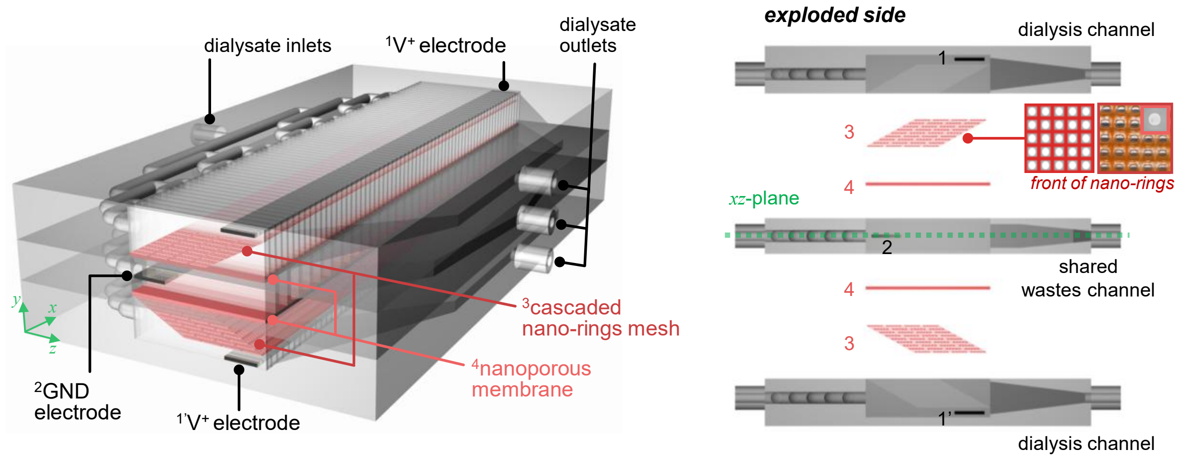


### **Supplementary Fig. 2.** Numerical simulation of cascaded nano-rings

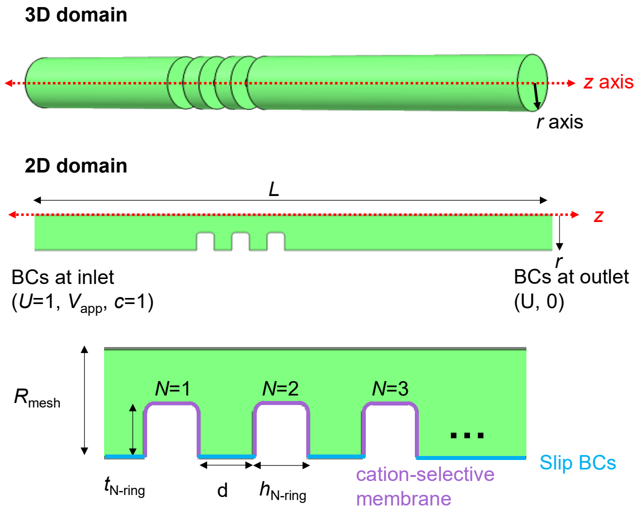


**Definition of parameters**

- **Applied voltage**: $V_{app}$ = 2
- **Inflow velocity**: $U$ = 1
- **Domain length**: $L$ = 15 (fix)
- **Radius of mesh**:

  $R_{mesh}$ = 1*$n$ ($n$ = [0.25, 0.5, 1, 2, 4, 8])
- **Thickness of coated-nafion**:

  $t_{N\text{-ring}}$ = $R_{mesh}$*$tp$ ($tp$ = [0.1, 0.2, …, 0.8])
- **Height of coated-nafion:** $h_{naf}$ = 0.25 (fix)
- **Distance of meshes**:

  $d$= $nd$*$h_{N\text{-ring}}$ ($nd$=[0.25, 0.5, 1, 2, 4, 8, 16, 32])
- **The number of meshes**: $N$=1, 2, 4, 8, 16

**Supplementary Fig. 3.** *In vitro* demonstration of cascaded nano-ring dialyzer

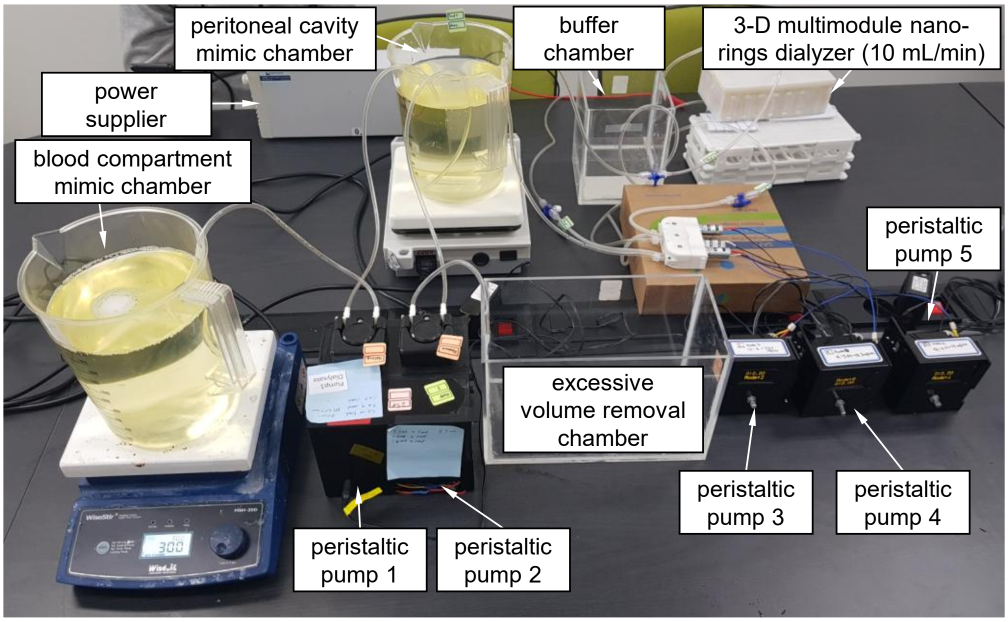


**Supplementary Fig. 4.** *In vivo* demonstration of cascaded nano-ring dialyzer

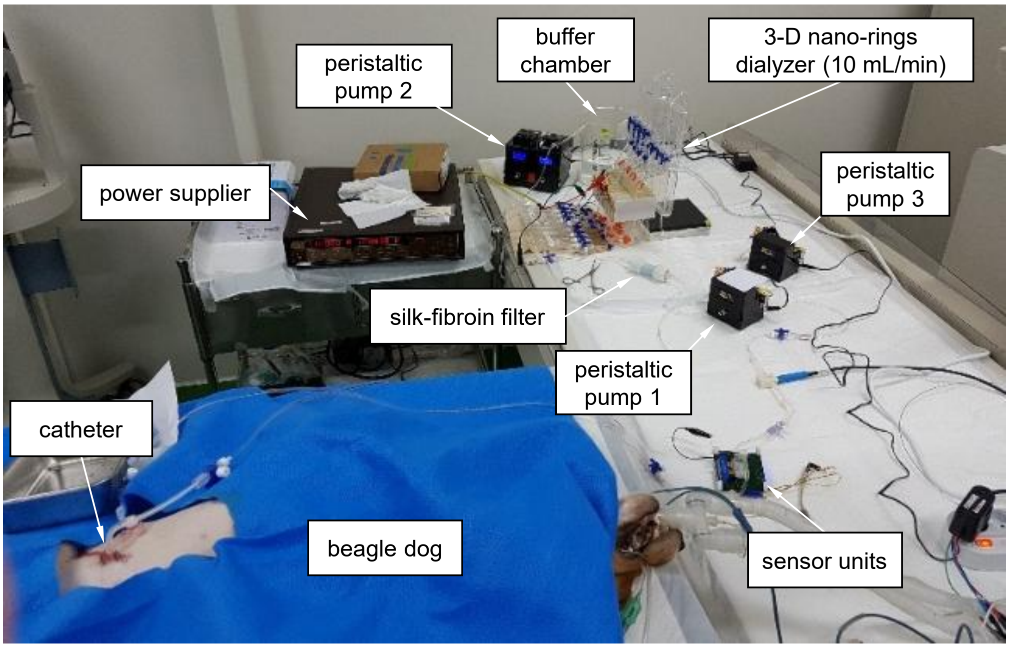

**Supplementary Fig. 5.** Block diagram for improved biocompatibility for prolonged continuous regenerative peritoneal dialysis

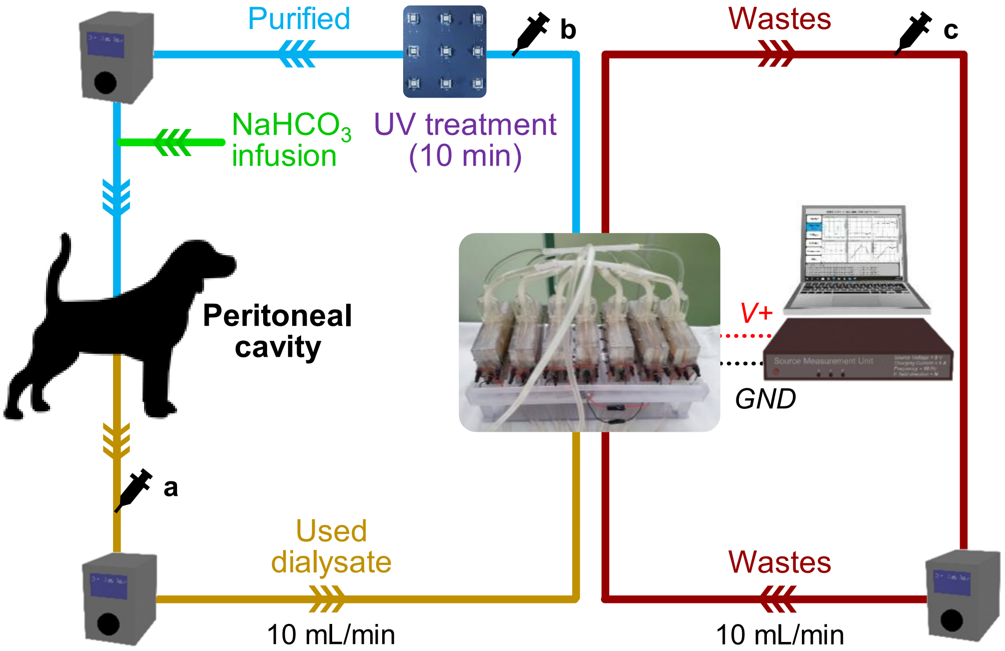


**Supplementary Fig. 6.** Integration of the nano-ring dialysis system into a portable bag-type configuration

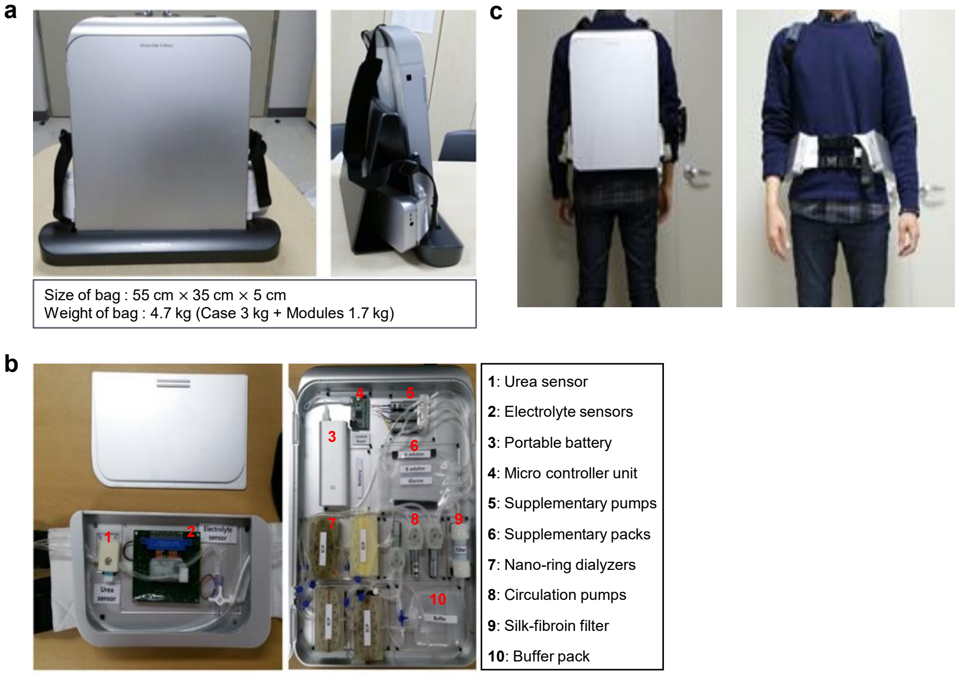
